# Neural Alpha Oscillations and Auditory Steady State Responses During Adaptation to a Cochlear Implant

**DOI:** 10.1101/2025.02.17.638598

**Authors:** Malte Wöstmann, Hannah Marie Meineke, Rainer Schönweiler, Daniela Hollfelder, Karl-Ludwig Bruchhage, Anke Leichtle, Jonas Obleser

**Affiliations:** Department of Psychology, University of Lübeck, Lübeck, Germany; Center of Brain, Behaviour, and Metabolism, University of Lübeck, Lübeck, Germany; Department of Clinical Research, University of Southern Denmark, Odense, Denmark; Department of Otorhinolaryngology, Phoniatrics and Paediatric Audiology, University Hospital of Schleswig-Holstein, Campus Lübeck, Lübeck, Germany; Department of Otorhinolaryngology, Head and Neck Surgery, University Hospital of Schleswig-Holstein, Campus Lübeck, Lübeck, Germany

**Keywords:** attention, auditory, cochlear implant, temporal coding, electroencephalography

## Abstract

The human auditory system must distinguish relevant sounds from noise. Severe hearing loss can be treated with cochlear implants (CIs), but how the brain adapts to electrical hearing remains unclear. This study examined adaptation to unilateral CI use in the first and seventh month after CI-activation using speech comprehension measures and electroencephalography (EEG) recordings, both during passive listening and an active spatial listening task. Neural phase-locking to amplitude-modulated (AM) sounds interacted with time, such that phase-locking longitudinally increased stronger for 40 compared with 4 Hz. In the spatial listening task, the benefit of performing the task with the CI on vs. off was most pronounced when the CI ear was primarily exposed to target speech. Lateralized alpha oscillations (∼10 Hz) reliably marked CI users’ focus of spatial attention. Stronger alpha modulation in the hemisphere opposite to the non-implanted ear indicates an attentional bias toward the acoustically hearing ear. Our findings suggest that adaptation to hearing with a CI is accomplished by dynamic changes in auditory phase-locking and a bias in auditory spatial attention.

**Data availability:** All data are available from the corresponding author upon request. A table including data for most important behavioural and neural outcome measures is available at https://osf.io/tkurz/.

## Introduction

Hearing loss is a common health condition, which affects a large group of individuals above the age of 60 years (> 65% according to WHO, 2021). In many cases, severe hearing loss can be treated with the implantation of a cochlear implant (CI), which transduces acoustic energy into electrical signals that stimulate the auditory nerve (Eshraghi et al., 2012). Unilateral CI users with combined electrical hearing on one ear through the CI and acoustic hearing on the contralateral ear pose a relevant test case for two reasons. First, there is a relatively large number of unilaterally implanted individuals, and we do at present not sufficiently well understand how the brain adapts to altered sensory perception following unilateral cochlear implantation. Second, models of auditory attention are based in large parts on findings in normal-hearing listeners with symmetrical hearing. Unilateral CI use offers a model for probing how a healthy, fully functioning neural system adapts to the unilateral restoration of degraded sensory input and its effects on auditory spatial attention allocation.

Successful hearing and communication rest on the interlocking of specific neural mechanisms. Accurate sound encoding and auditory perception aid auditory object formation, whereas the allocation of auditory attention to enhance neural representations of target sounds and to suppress noise aids auditory object selection (Shinn-Cunningham, 2008). In CI users, listening performance on average increases during the first six months after cochlear implantation, followed by a plateau (e.g., Lenarz et al., 2012). With the present study, we aim to investigate the neural dynamics of auditory perception and spatial attention in unilateral CI users during the first half year following cochlear implant activation.

Sound processing through a CI primarily degrades spectral cues but leaves temporal envelope cues largely intact. Thus, CI users are thought to rely mainly on temporal-envelope information (Dincer D’Alessandro et al., 2018; Heng et al., 2011; Rosen, 1992; Shannon et al., 1995). Here, we ask how adaptation to hearing with a CI reflects in neural phase-locking to the sound envelope. In the human electroencephalogram (EEG) the auditory steady-state response (ASSR), also referred to as the envelope following response (EFR) in case of a periodic envelope, has been linked to auditory temporal processing acuity (Purcell et al., 2004; Roß et al., 2002). ASSRs evoked by 40-Hz modulation have a particularly high SNR (Galambos et al., 1981) and are generated in auditory cortical and subcortical regions (David et al., 2023; Farahani et al., 2017; Luke et al., 2017), whereas ASSRs evoked by slower frequencies (<20 Hz) originate primarily from auditory cortex (Liégeois-Chauvel et al., 2004). The 40-Hz ASSR is altered in central brain disorders (e.g., Grent-’t-Jong et al., 2023; Kwon et al., 1999; Wilson et al., 2007) and has been associated with cortical inhibition (e.g., Toso et al., 2024).

In CI users, 4- and 40-Hz ASSRs relate to modulation detection thresholds, indicating their feasibility to assess temporal coding in CI users (Luke et al., 2015). Somewhat unintuitively, larger cortical representations of slow amplitude modulations (∼4 Hz) have been observed at older age (Anderson et al., 2020; Presacco et al., 2016) and in hearing loss (Fuglsang et al., 2020; Orf et al., 2025), which might suggest an imbalance of excitatory and inhibitory processing. Here, we test specifically how adaptation to hearing with a CI reflects in changes of the 4-versus 40-Hz ASSR.

A prominent neural signature of spatial attention in different sensory modalities is the hemispheric lateralization of ∼10 Hz alpha oscillatory power (auditory: Ahveninen et al., 2013; somatosensory: Haegens et al., 2011; visual: Worden et al., 2000). In normal-hearing listeners, alpha power increases in the hemisphere ipsilateral to the focus of auditory attention and decreases in the contralateral hemisphere (e.g., Deng et al., 2020; Müller & Weisz, 2012; Wöstmann et al., 2016). Relatively reduced versus enhanced alpha power is thought to reflect the attentional selection of targets and suppression of distraction, respectively (e.g., Bonnefond & Jensen, 2024; Schneider et al., 2021; Strauß et al., 2014). Through experimental separation of lateral target versus distractor processing, we have recently shown that there are two lateralized alpha responses (Wöstmann et al., 2019), one related to the selection of targets and another to the suppression of distraction (for related findings in the visual modality, see Cruz et al., 2025; Yang et al., 2024).

Evidence for auditory attentional alpha power modulation in CI users is scarce, however. Alpha power modulation in CI users has been found to relate to the subjective experience of task difficulty, i.e., listening effort (Dimitrijevic et al., 2019). Furthermore, in bilateral CI users, Paul and colleagues (2020) found evidence for alpha lateralization during auditory spatial attention to speech. However, we do at present not understand how asymmetric hearing loss and adaptation to hearing with a unilateral CI affect large-scale neural network organization (see also Jiwani et al., 2021) and the allocation of auditory spatial attention.

Here, we present data from a longitudinal EEG study in unilateral CI users to probe neural signatures of auditory perceptual processing and spatial attention allocation in the first and seventh month following CI-activation. We asked, first, whether the relative pattern of slow (4-Hz) versus fast (40-Hz) ASSR responses would reflect a change in the excitation-inhibition balance during adaptation to listening with a CI. Second, we tested whether unilateral CI users exhibit asymmetry in the neural allocation of spatial attention depending on the side of the CI.

## Methods

### Participants

N=20 native German speakers were recruited for this study. Two participants dropped out before finishing the first recording session. Three participants dropped out of the study after completing the first session. Part of the EEG data of one participant were not recorded due to technical issues. Thus, N = 18 participants with complete data of the first session and N=14 complete datasets with two recording sessions were available for analysis. Of these N = 14 participants with complete datasets, all but one participant were right-handed according to the Edinburgh handedness questionnaire (Oldfield, 1971).

All participants were unilaterally fitted with a cochlear implant (CI) at the Department of Otorhinolaryngology, Head and Neck Surgery, University Hospital of Schleswig-Holstein, Campus Lübeck, Germany, with a CI from the manufacturers Cochlear or MedEL. Their contralateral hearing loss varied from normal hearing to profoundly hard of hearing (see also Fig. 2B). Participants were financially compensated for their participation at an hourly rate of €10. All participants gave written informed consent to participate in the study. All procedures were approved by the ethics committee of the University of Lübeck, Lübeck, Germany (ID of approval: 19–127).

### Study timeline and setup

Participants completed the same test battery twice; once approximately in the first month after activation of the CI (Session 1) and once approximately six months thereafter (Session 2). Each session comprised two data recordings on two separate days. Time intervals provided below refer to the second data recording of each session, which included the main tasks reported here (i.e., passive and active listening tasks in the EEG). For the N=14 complete datasets, the average time interval between CI-activation and Session 1 was 3.3 weeks (range: 1.3–7.3). The average interval between Session 1 and 2 was 6,4 months (range: 5.5– 7.8).

Data recording spanned the time window from September 2020 to May 2023. Sound stimuli were presented via loudspeakers (Genelec 8020D), placed at a distance of ∼70cm to the participant’s head. Depending on the task and experimental condition, loudspeakers were positioned on the left side (–90°), right side (+90°) or in the front (0°). Individual tests (AMRD, passive listening, spatial attention task) were implemented in Matlab (R2017b) and Psychtoolbox (Brainard, 1997).

In addition to the individual tests of the test battery explained below, participants performed a continuous speech tracking task (adapted from Kraus et al., 2021), which was not analysed for the purpose of the present study.

### Audiological tests

Pure tone audiograms (at frequencies 0.125, 0.25, 0.5, 1, 2, 3, 4, 6, and 8 kHz) and speech intelligibility scores in the “Freiburg Monosyllabic Speech Test” (FMST, 65 dB SPL) were collected prior to CI implantation and at the regular clinical appointment of the participant. The audiological measurements were conducted for both ears separately, if possible. The pre-operative audiogram and FMST was measured unaided, the post-operative measurements were conducted aided with the CI. The contralateral ear measurements were conducted aided with a hearing aid when applicable.

Temporal sensitivity was assessed with an adaptive amplitude modulation rate discrimination (AMRD) test (for details, see Erb et al., 2019). In a Three-Alternative Forced Choice (3-AFC) adaptive staircase paradigm, three broadband noise sounds were presented in sequence on each trial. Two standard stimuli were amplitude-modulated at 4 Hz and one deviant stimulus was amplitude-modulated at a frequency of 4–6 Hz (determined according to the adaptive staircase method). The task always started with a 6-Hz deviant. The position of the deviant varied randomly on a trial-by-trial basis. Participants had the task to report the position (1–3) of the deviant via button press. A 2-down, 1-up adaptive staircase procedure determined the AMRD threshold converging on 70.7% correct responses (Levitt, 1971). The initial step size was 0.5 Hz, which was reduced to the minimal step size of 0.25 Hz after four reversals. The test terminated after 10 reversals. The AMRD threshold corresponded to the average AM rate difference of the deviant vs. standard across the last six reversals. Participants completed the AMRD task twice; the minimum of the two resulting thresholds was used for further analyses.

Participants filled out the short form of the “Speech, Spatial and Quality of Hearing Scale” (SSQ, German short version; Kießling et al., 2011) at the beginning of each of the two sessions to assess their subjective hearing performance (Gatehouse & Noble, 2004). As an outcome measure, we used the sum of scores across all items, with a higher score corresponding to better self-rated hearing.

### Threshold estimation procedure

As absolute hearing levels differ dramatically between CI users, we aimed to present sound stimuli at an overall intensity that was adapted to individual hearing thresholds. To this end, absolute hearing thresholds were assessed using the method of limits. We used the same spoken numbers (1-9, female voice) as for the spatial attention task. For each speaker location (front, left, right) spoken numbers were presented in random order, starting at an inaudible intensity and increasing in steps of 1dB. Participants were instructed to press a button as soon as they could hear a number. This procedure was repeated three times. Afterwards, three repetitions of the same procedure followed with the exception that presentation started far above threshold and decreased in steps on 1dB. Participants were instructed to press a button as soon as they could not hear the numbers anymore. The threshold was determined by averaging across scores of the last two repetitions for increasing and the last two repetitions for decreasing sound intensity. Thresholds for the front speaker location were always measured first, followed by the non-CI side and CI side. Thresholds varied considerably across participants and conditions (range: 43.75 dB), but a repeated-measures ANOVA revealed no main or interaction effects of Speaker location (front, left, right) or Session (1 vs. 2) on thresholds (all *p* > .33).

Determined thresholds were considered to correspond to 0dB sensation level (SL) and sound stimuli in the passive listening task and spatial attention task were presented at +40dB SL at each individual loudspeaker position. Separate thresholds measurements were conducted in Session 1 and Session 2. Alternatively, it would have been possible to use participants’ pure tone thresholds for individual stimulus adjustments, especially for the passive listening task with amplitude-modulated sounds. However, since thresholds estimated from our procedure were highly correlated with pure tone thresholds and since the assessed auditory steady-state response (ASSR) in the passive listening task did not significantly relate to pure tone thresholds (see Fig. S3), there is no indication of bias in our threshold estimation procedure.

### Passive listening to amplitude-modulated (AM) sounds

Participants listened passively to 2-s long 1000-Hz tones, with three different AM rates: 4, 20, & 40 Hz (100% modulation depth). Sounds were presented binaurally at +40dB SL over the left and right loudspeaker. In total, 180 sounds were presented (60 per AM rate) in a pseudo-randomized order, making sure no more than three consecutive sounds would have the same AM rate. The inter-stimulus interval was randomly jittered in 100-ms steps between 500 and 1500ms.

Of note, the 20-Hz AM rate was not of primary interest in the analysis of the auditory steady-state response (ASSR). Instead, this condition was used in the experiment to increase the ability to separate auditory neural components (which should exhibit stronger phase-locking to 4 & 40 Hz compared with 20 Hz) from components related to the processing in the cochlear implant (which should exhibit similar phase-locking across frequencies).

### Active listening in an auditory spatial attention task

The design of the auditory spatial attention task was adapted from Wöstmann et al., (2019). The overarching aim was to implement listening conditions that separate attentional selection from suppression of lateralized sounds on the CI-versus non-CI-side. To this end, two competing spoken numbers were presented from two loudspeaker positions. One loudspeaker was always positioned in the front (0°), while the other changed its position from left (–90°) to right (+90°) in a block-wise fashion. This way, the following four conditions were implemented: *select CI-side, select non-CI-side, suppress CI-side, suppress non-CI-side*.

On each trial, participants were first presented for 500 ms with a spatial cue (^, <, or >) to indicate the location of the upcoming target sound (i.e., front, left, or right). In the ensuing anticipation period, randomly jittered in duration between 1000 and 1800 ms in steps of 100 ms, a fixation cross was presented on the screen. Then, two different, randomly selected numbers were presented, each one over one of the two loudspeakers. The set of numbers (adapted from Obleser et al., 2012) comprised the numbers 1–9, which were spoken by a female voice and adjusted to a duration of 500 ms each. To enhance the perception of simultaneous onsets of the two numbers on each trial, we applied a perceptual onset matching (as described in Wöstmann et al., 2016). Participants used a numpad to indicate the target number in the end of each trial.

The task comprised six blocks. Loudspeakers were positioned in the front and on the left in half of the blocks and in the front and on the right in the other half. Each block contained 50 trials, 25 with an attention cue to the front and 25 with a cue to the side (left or right). The loudspeaker setup (front & left versus front & right) alternated from block to block, starting with the front & left setup for participants with the CI on the right side and vice versa for participants with the CI on the left side. In total, the task contained 300 trials with 75 trials per experimental condition. During each block, continuous broadband background noise (band pass filtered between 0.1 and 8 kHz) was presented from an additional loudspeaker positioned behind the participant at a low intensity of +25 dB (SL; as determined for the front loudspeaker in the threshold estimation procedure).

The task was implemented as an adaptive 1-down, 1-up staircase procedure (Levitt, 1971) targeting the 50% speech reception threshold (SRT_50_). For each of the four experimental conditions, the task started at an SNR of +10dB (except for the first participant in the first session, for whom it started at 0 dB), meaning that the target intensity was 10 dB higher than the distractor intensity. Following a correct response in one condition, the SNR was lowered by 2dB for the next trial in this condition. Following an incorrect response in one condition, the SNR was increased by 2dB for the next trial in this condition. The threshold in each condition was calculated by averaging the SNR for all trials except for the first block in which this condition was implemented. In Session 1, three participants had partially missing behavioural data (last two blocks in CI off recording missing for ID 4&5; last block in CI off recording missing for ID 7).

In each session (1&2), each participant performed the auditory spatial attention task twice, first with the CI switched on and then, after a short break, with the CI switched off.

### EEG preprocessing and CI artifact detection

The EEG was recorded at 64 active scalp electrodes (Ag/Ag-Cl; ActiChamp, Brain Products) at a sampling rate of 1000 Hz, with a DC–280 Hz bandwidth, against reference electrode Fz. All electrode impedances were kept below ∼30 kΩ. Note that depending on the placement of the cochlear implant (and the hearing aid on the contralateral side) some electrodes above the device could not be connected and were interpolated later (see below). To ensure equivalent placement of the EEG cap, the vertex electrode (Cz) was placed at 50% of the distance between inion and nasion and between left and right ear lobes. For EEG data analysis, we used the FieldTrip toolbox (Oostenveld et al., 2010) for MATLAB (R2013b/R2023b) and custom script.

We used the data from the passive listening task to identify independent components in the EEG data relating to artifacts, auditory activity, and other brain activity for further analyses. To this end, the continuous data were filtered (high pass: 0.1 Hz, low pass: 100 Hz) and split in epochs around sound onsets (–.5 to 2.5 s). Next, trials for which any channel exceeded a range of 300 μV in the time interval 0–2 s were removed. An independent component analysis (ICA) was used to split the data into a number of independent components required to explain 90% variance. Component time courses, topographic maps, and frequency spectra were carefully inspected (see Fig. 1) to identify (i) components relating to the CI artifact (sharp, low-latency sound-evoked activity, largest at electrodes close to CI), (ii) components relating to auditory neural processing (sound-evoked pattern of P1-N1-P2 components, strongest at central electrodes), (iii) other brain components (components with typical characteristics of electrophysiological brain activity, such as 1/f spectra), and (iv) remaining artefactual components (relating e.g., to eye-blinks or saccades, muscle activity, or line noise). Due to the characteristic differences of components related to CI processing versus auditory neural activity in topography and latency, CI-artefact removal from the EEG data was possible in the present dataset.

**Figure 1.**
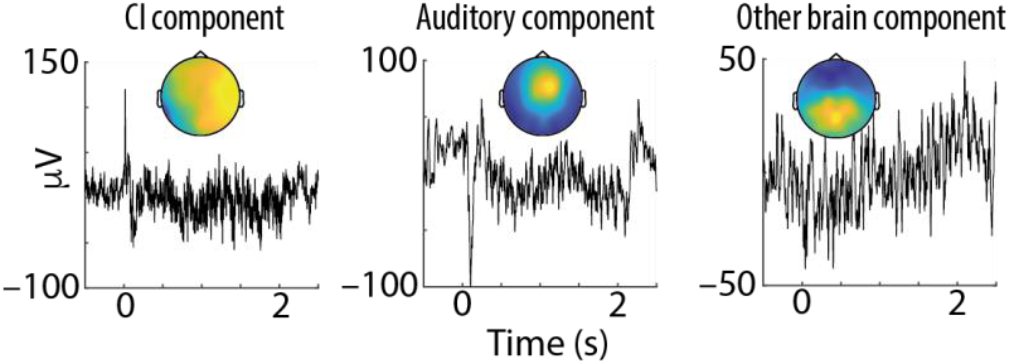
Time courses (averaged across all trials and AM rates in the passive listening task) and topographic maps of exemplary ICA components for one participant with the CI on the right side (ID 5, session 1). The CI artifact component shows a sharp, low-latency response to sound onset and largest weights on the CI side. The auditory component has highest weights at central electrode sites and shows typical auditory onset and offset responses in the beginning (0 s) and end (2 s) of sound presentation, respectively. The ‘other brain component’ depicted here has high weights at parietal electrodes, as it is typical for alpha (∼10 Hz) oscillatory power.

**Figure 2.**
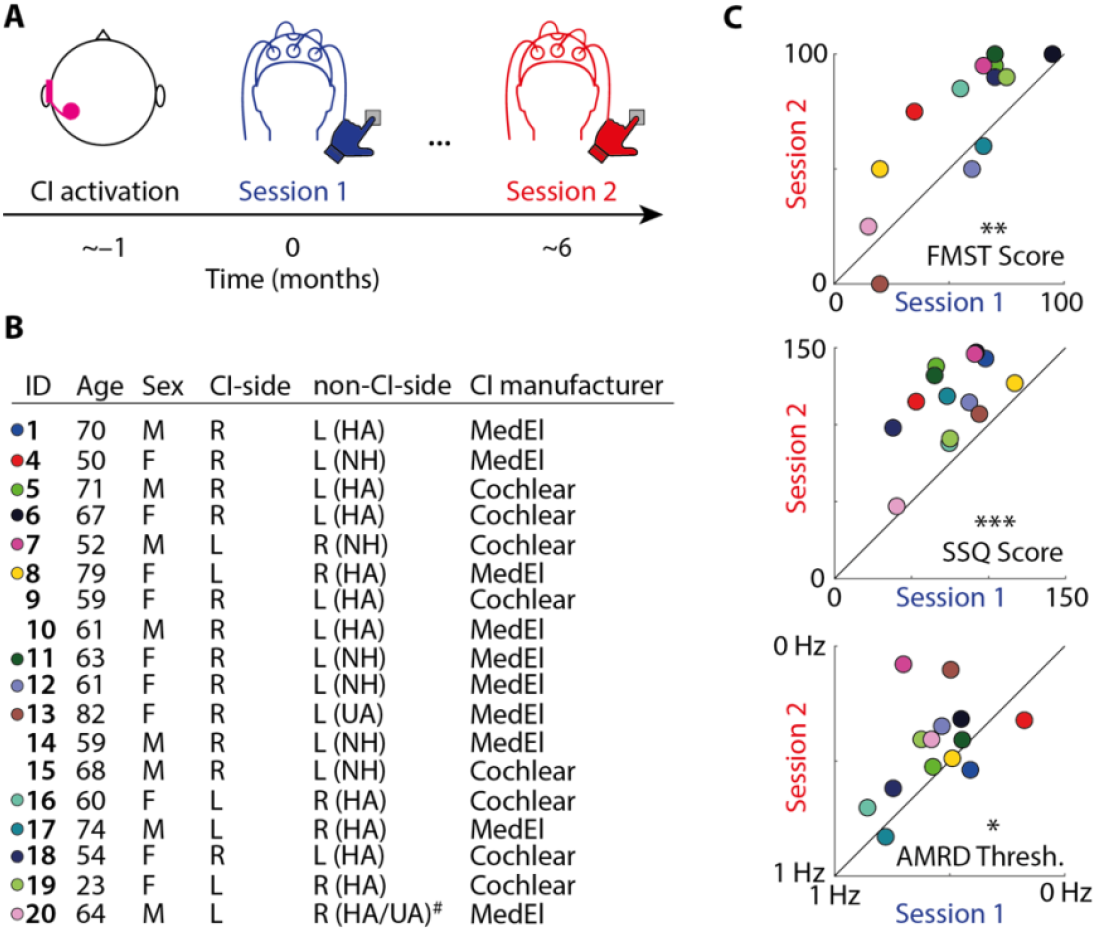
(**A**) Schematic depiction of the general study design. Unilateral CI users completed behavioural and electrophysiological measurement sessions in the 1^st^ month (Session 1) and 7^th^ month (Session 2) after CI activation. (**B**) Participant information. Age refers to the timepoint of the first session. R: right, L: left, HA: hearing aid, NH: normal hearing (no or mild hearing loss according to WHO criteria), UA: unaided. ^#^ Participant had significant deterioration of hearing on non-CI side in-between Session 1 & 2 and could not be adequately fitted with a hearing aid (HA in Session 1; UA in Session 2). Coloured circles indicate individual participants who completed both sessions (S1 & S2) and had no missing data (N = 14; the same colour-coding is used in all forthcoming figures). (**C**) 45-degree plots show percentage correct scores in the ‘Freiburg monosyllabic speech test’ (FMST; top), overall score in the ‘speech, spatial and qualities questionnaire’ (SSQ, summed over items from all three scales; middle), and thresholds in the ‘amplitude modulation rate discrimination task’ (AMRD; bottom). Note that x- and y-axes for the AMRD results are reversed, such that data points above the diagonal indicate improved (i.e., lower) thresholds for Session 2 vs. 1. * *p* < .05, ** *p* < .01, *** *p* < .001.

### Analysis of EEG data in the passive listening task

Continuous EEG data of the passive listening task were filtered (high pass: 0.1 Hz, low pass: 100 Hz) and split in epochs around sound onsets (–.5 to 2.5 s). To focus on auditory activity, we projected the data though the previously identified auditory components from each participant (separately for data in Session 1 and 2). Next, missing channels were interpolated (spline interpolation) and the data were re-referenced to the average of all channels. Epochs with 10% largest ranges (in time interval 0–2 s; at any channel) were removed.

To quantify the magnitude of the neural onset response to amplitude modulated sound stimuli, we quantified global field power (GFP) separately for each Session (1, 2), participant, and AM rate (4, 20, 40 Hz) by calculating the standard deviation of EEG activity across electrodes in the time interval spanning obligatory onset responses (P1, N1, & P2; 0–230 ms).

To analyse the auditory steady-state response (ASSR), epochs were split according to the three AM rates (4, 20, 40 Hz) and the event-related potential (ERP) was calculated by averaging over trials in the time domain. To focus on activity unrelated to the sound onset-evoked response, we cut out ERPs in the time interval 0.5–2 s and then obtained spectral power using Fast Fourier Transform (FFT; rectangular window) for frequencies 1–60 Hz in steps of 1 Hz with 2 Hz spectral smoothing. Finally, spectral power was averaged over nine fronto-central electrodes (Fz, F1, F2, FCz, FC1, FC2, Cz, C1, C2).

### Analysis of EEG data in the spatial attention task

Continuous EEG data of the auditory spatial attention task were filtered (high pass: 1 Hz, low pass: 100 Hz) and split in epochs around visual cue onset (–1 to 4 s). We used the previously identified ICA components (see above), projected the data though these components, and rejected CI and other artefactual components (separately for data in Session 1 and 2). Missing channels were interpolated (spline interpolation) and the data were re-referenced to the average of all channels. Epochs with 10% largest ranges (in time interval 0–2.5 s; at any channel) were removed. Finally, epochs were split into experimental conditions. For three participants in session 1 (ID 4, 5, 16), one block of EEG data was missing and thus not included in the analysis.

Based on previous studies in healthy, normal hearing participants (e.g., Wöstmann et al., 2016, 2019), we focused the spectral analysis of oscillatory power on the anticipation period in the cue-target interval. To this end, we cut out the EEG data in the time interval 0.75 to 1.5 s, which is temporally remote from cue-evoked activity and from any sound-induced activity (>1.5 s). Spectral power was obtained using Fast Fourier Transform (FFT) with multi-tapering (DPSS, discrete prolate spheroidal sequences) for frequencies 1–30 Hz in steps of 1 Hz with 2 Hz spectral smoothing.

Lateralization of spectral power was calculated individually for each session (1, 2), CI status (on, off), and participant for the selection of lateral targets [LI_selection_ = (Pow_select-left_ – Pow_select-right_) / (Pow_select-left_ + Pow_select-right_) and for the suppression of lateral distractors [LI_suppression_ = (Pow_suppress-left_ – Pow_suppress-right_) / (Pow_suppress-left_ + Pow_suppress-right_).

For further analyses, we followed in part a preregistered analysis plan for a previous study (https://osf.io/bv7zs; Wöstmann et al., 2019). We averaged each lateralization index (LI_selection,_ LI_suppression_) across frequencies in the alpha band (7 –13 Hz), separately for two sets of 12 left and 12 right hemispheric occipito-parietal electrodes (TP9/10, TP7/8, CP5/6, CP3/4, CP1/2, P7/8, P5/6, P3/4, P1/2, PO7/8, PO3/4, and O1/2).

### Statistical analyses

To estimate the reliability of audiological tests (FMST, SSQ, AMRD) from results in Sessions 1 vs. 2, we used Spearman correlation coefficients. Nonparametric permutation tests were used to test for mean test-score differences in Session 1 vs. 2. The reported p-value (denoted *p*_perm_) corresponds to the proportion of absolute values of 10,000 dependent-samples *t*-statistics computed on data with permuted condition labels exceeding the absolute empirical *t*-value for the original data.

For the EEG outcome measures in the passive listening task (GFP & ASSR), we used repeated-measures ANOVAs with the factors Session (1 vs. 2) and AM rate of interest (4 vs. 40 Hz).

For behavioural data in the spatial attention task, SRT_50_ values for individual conditions were determined and CI benefit was calculated by subtracting the SRT_50_ for CI on – CI off. CI benefits were then submitted to a repeated-measures ANOVA (including Greenhouse-Geisser correction of degrees of freedom to control for violation of sphericity) with the factors Session (1 vs. 2) and Condition (select CI-side, suppress CI-side, select non-CI-side, suppress non-CI-side), followed by post-hoc nonparametric permutation tests. Only complete datasets (recordings in Session 1 & 2) were included in this analysis. Three participants with complete datasets were excluded (ID 6, 18, 20), as they had unusually high SRTs (> +40 dB) in at least one condition, leaving N = 11 participants with complete datasets for the analysis of behaviour in the spatial attention task.

For EEG data in the spatial attention task, we first tested for significant hemispheric modulation of LI_selection_ and LI_suppression_. To this end, we used two repeated-measures ANOVAs on average alpha (7–13 Hz) power across 12 left and 12 right hemispheric occipito-parietal electrodes (see above) with the factors Session (1 vs. 2), CI status (on vs. off) and Hemisphere (left vs. right). Next, we tested whether the significant hemispheric modulation of LI_selection_ would be stronger in the hemisphere on the same versus opposite side of the CI. For each participant, we calculated two alpha lateralization indices (ALI; Wöstmann et al., 2016), one contrasting selection of ipsi-versus contralateral targets with respect to electrodes on the CI-side (ALI_CI-side_) and the other with respect to electrodes on the non-CI-side (ALI_non-CI-side_). A repeated-measures ANOVA with the factors Session (1 vs. 2), CI status (on vs. off) and Hemisphere (CI-side vs. non-CI-side) was used for statistical testing.

## Results

Here, we employed an audiological and neuro-behavioural test battery in unilateral CI users at two time points; in the first and seventh month following CI activation (Fig. 2A). We hypothesized that the to-be-expected improvements in subjective and objective hearing abilities following CI activation would be accompanied by improved neural perceptual processing of sound. Furthermore, we expected that asymmetries in the neural implementation of auditory spatial attention would be explained by the side of the CI.

### Improvements in subjective and objective hearing

We administered three tests to assess speech comprehension (FMST), subjective hearing abilities across different domains (SSQ), as well as auditory temporal discrimination ability (AMRD). To probe whether the tests are reliable, we calculated test-retest reliability, which was moderate to high (Spearman’s rho; FMST: *rho* = 0.819; SSQ: *rho* = 0.442; AMRD: *rho* = 0.392). Compared with Session 1, non-parametric permutation tests revealed significant improvements in Session 2 for all tests (Fig. 2C; FMST: *p*_*perm*_ = 0.007, average performance increase = 15.36%; SSQ: *p*_*perm*_ < 0.001, average improvement = 39.11 scores; AMRD: *p*_*perm*_ = 0.026, average threshold decrease = 0.126 Hz).

### Passive listening: Differential modulation of low-vs. high-frequency phase locking

CI users listened passively to amplitude-modulated (AM) 1000-Hz tones (Fig. 3A). Most importantly, we used AM rates of 4 vs. 40 Hz, which are thought to reveal the neural tracking of slow modulations as they occur in spoken language vs. temporal fidelity in tracking fast modulations, respectively. Additionally, we included a 20 Hz AM modulation condition.

**Figure 3.**
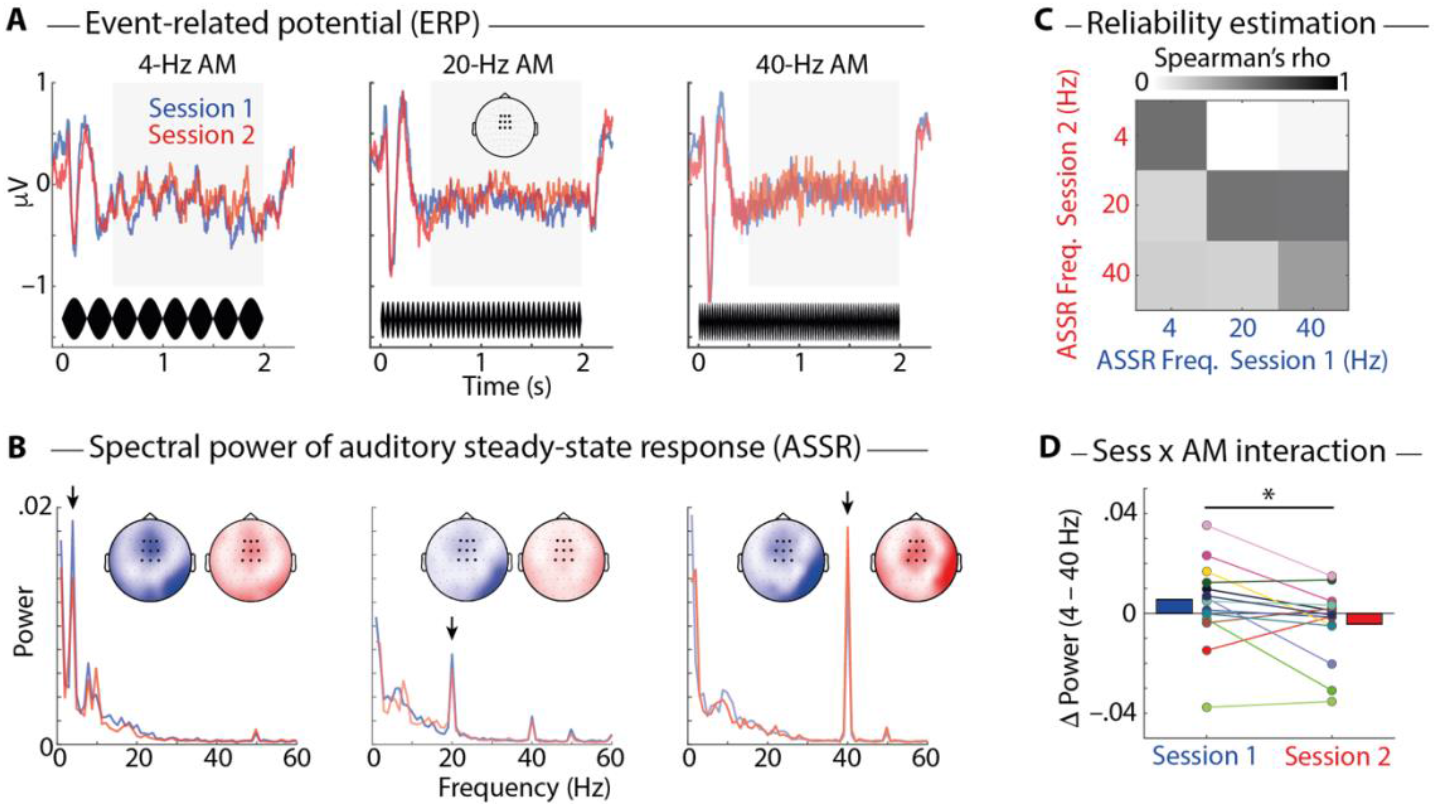
(**A**) Event-Related Potential (ERP), averaged across participants in Session 1 (N = 18, blue) and in Session 2 (N = 14, red) and 9 fronto-central channels (highlighted in topographic map). Amplitude-modulated (AM) sound stimuli are shown in black in the bottom. (**B**) Lines show average spectral power, calculated on the ERP in the time-interval 0.5–2s (grey boxes in A), averaged across 9 fronto-central channels. Arrows highlight corresponding AM rates at 4, 20, and 40 Hz. Topographic maps show spectral power (color bar limits: 0–0.04) at corresponding AM rates for Session 1 (blue) and 2 (red). Single-subject ERPs and spectral power data are shown in Fig. S1 & S2. (**C**) Correlation matrix shows Spearman’s rho for ASSR peak frequencies correlated between Session 1 & 2 (N = 14). (**D**) Visualization of significant Session (1 vs 2) x AM rate (4 vs 40 Hz) interaction. Bars show average spectral power, colored lines and dots show individual participants. * *p* < 0.05.

The passive listening task served two purposes. First, an independent component analysis (ICA) could be used to identify EEG components related to the CI artifact, to auditory activity, and to other brain activity. A selection of these components could then be used for subsequent analyses. Second, we rejected all but the auditory components from these data and calculated the auditory steady-state response (ASSR) in the EEG to probe CI users’ auditory perceptual processing.

The Event-Related Potential (ERP; Fig. 3A) to AM sounds showed obligatory onset response components (P1, N1, P2), followed by a steady-state response – the ASSR – corresponding to the AM rate. The onset response appeared to increase for higher AM rates. For statistical analysis, we quantified the overall onset response by Global Field Power (GFP; i.e., the standard deviation across electrodes) in the time interval 0–230 ms following sound onset. There was a significant main effect of AM rate (4 vs. 40 Hz) on GFP (*F*_1, 13_ = 5.367; *p* = 0.037), indicating larger GFP for the 40-Hz compared with the 4-Hz condition (*p*_*perm*_ = 0.029). There was neither a main effect of Session nor a Session x AM rate interaction (both *p* > 0.12).

Spectral power of the ASSR was calculated in the time interval 0.5–2 s following sound onset. Prominent spectral peaks corresponding to AM rates of sounds were present in these spectra (Fig. 3B). Spectral power of ASSR peak frequencies was moderately correlated across the two sessions and generally higher for corresponding frequencies (Fig. 3C), which demonstrates reliability of the ASSR measure. Critically, for the two AM rates of interest (4 & 40 Hz), there was a significant Session x AM rate interaction (Fig. 3D; *F*_1, 13_ = 6.066; *p* = 0.029). Post-hoc tests revealed that for individual frequencies, there were no significant effects of Session (4 Hz: *p*_*perm*_ = 0.127; 40 Hz: *p*_*perm*_ = 0.459). The significant interaction was nevertheless robust and persisted when the spectral analysis was extended to incorporate the entire stimulus duration (0–2s; *F*_1, 13_ = 5.380; *p* = 0.037), as well as when regressing out differences in threshold estimates (which were used to individualize stimulus presentation levels) on the CI-side (*p* = 0.0345) and non-CI-side (*p* = 0.0286).

Finally, we asked whether the observed AM rate (4 vs. 40 Hz) x Session (1 vs. 2) interaction was driven by a change in band-specific EEG amplitude or phase-consistency, since both reflect in the spectral power of the ASSR. However, when analysing inter-trial phase coherence (ITPC; which is in theory independent of spectral power) this effect was likely absent (*F*_1, 13_ = 0.608; *p* = 0.450; Bayes Factor, *BF*_10_ = 0.398), suggesting that the AM rate effect is not solely driven by modulation of phase coherence.

### Active listening: Hearing with a single-side CI benefits spatial listening

To investigate the benefits of listening with a unilateral CI in different spatial arrangements of competing target versus distractor speech, participants performed a spatial listening task with the CI turned on versus off (Fig. 4 A&B). Task performance was continuously titrated using an adaptive staircase procedure, such that the SNR was adjusted for each task condition separately to arrive at 50% task accuracy (Fig. 4C). In each run of the task, participants performed six blocks, wherein three blocks presented one sound source on the left and three on the right (while the other sound source was always presented in the front). The speech reception threshold (SRT_50_) for a specific condition corresponded to the average SNR across all trials of the respective condition, except for trials in the first block. Participants performed two runs of this task, first with the CI switched on and thereafter with the CI switched off.

**Figure 4.**
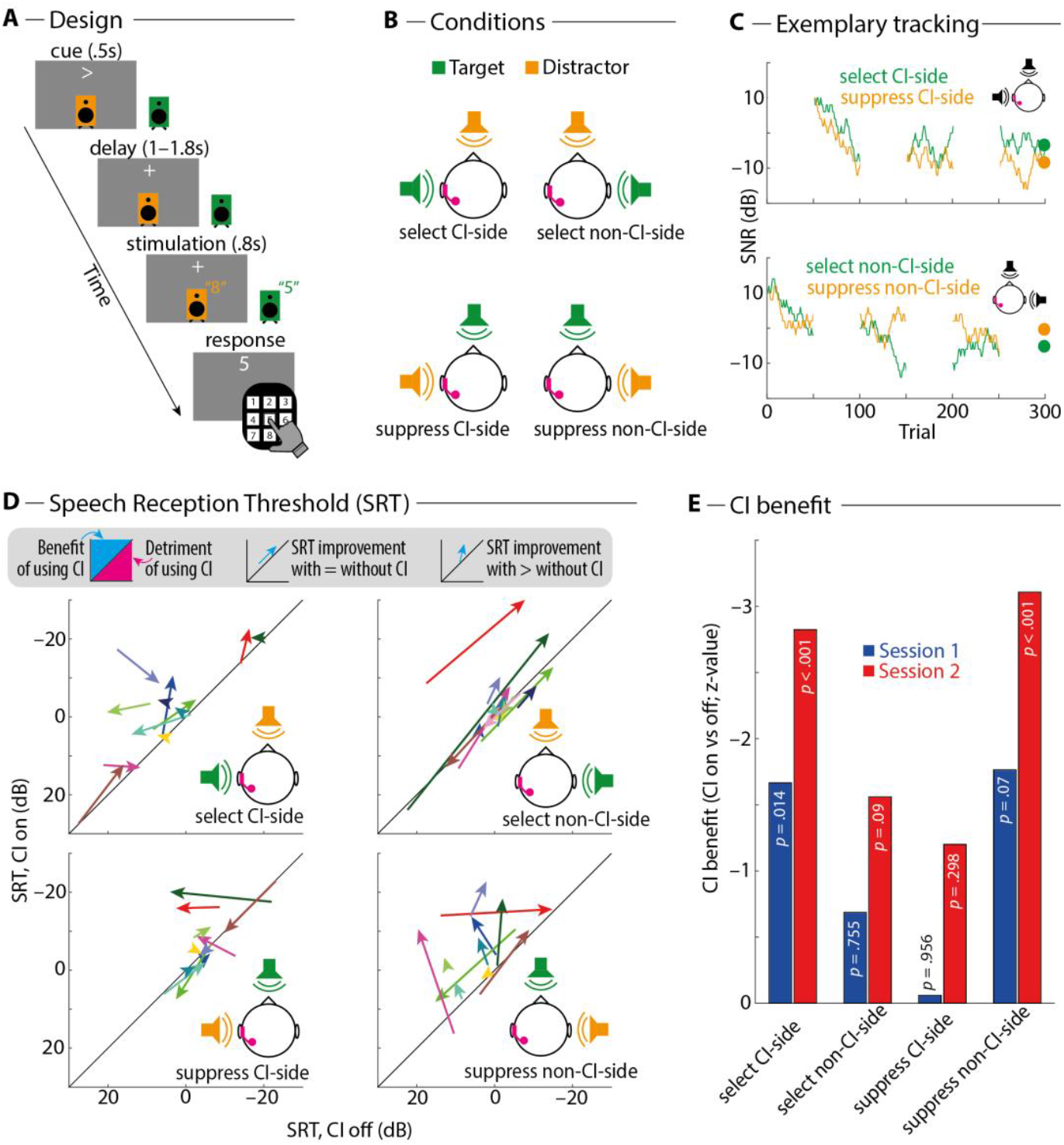
(**A**) Design of spatial listening task. On each trial, one of two loudspeakers was the target (green; indicated by visual cue) and the other was the distractor (orange). The task was to attend to and report the spoken number presented over the target loudspeaker. (**B**) Experimental conditions for an exemplary participant with the CI on the left side. (**C**) The SNR was continuously titrated throughout the task to achieve 50% task accuracy in each condition. Graphs show the adaptively tracked SNR over time for one exemplary participant (ID 19, Session 1). Note that the speaker setup (front & left versus front & right) alternated in a block-wise fashion, meaning that only two of the four conditions were presented (and thus titrated) in a respective block. Large dots in the end of the tracking procedure indicate the speech reception threshold (SRT_50_), which was calculated as the average SNR across all trials of the respective condition, except for trials in the first block. (**D**) Arrows show individual participants’ SRT for the CI switched off (x-axis) versus on (y-axis), pointing from the respective SRT in Session 1 to Session 2 (N = 11). More negative (i.e., better) SRTs are plotted upward (y-axis) and to the right (x-axis). Data points above the diagonal indicate better SRT when the CI was switched on versus off (i.e., CI benefit). (**E**) Summary and statistical analysis of the data shown in (D). Bars show z-values for the statistical contrast CI on versus off for individual task conditions and sessions. Negative z-values (corresponding to more negative – i.e., better – SRT when the CI was switched on) are plotted upwards. P-values in bars correspond to results of non-parametric permutation tests of CI benefits against zero.

Overall, SRTs were lower (i.e., better) with the CI switched on versus off (Fig. 4 D&E), which speaks to a CI benefit. Statistical analysis revealed a significant main effect of Condition on the CI benefit (*F*_1.63, 16.28_ = 4.415; *p* = 0.036), but the main effect of Session (*F*_1, 10_ = 3.722; *p* = 0.083) and the interaction of the two were not statistically significant (*F*_2, 19.97_ = 0.543; *p* = 0.589). Post-hoc nonparametric permutation tests showed that a significant benefit of the CI was only observed in conditions where the implanted ear was primarily exposed to the target sound (select CI-side: Session 1, *p*_*perm*_ = 0.014; Session *2, p*_*perm*_ < 0.001; suppress non-CI-side: Session 1, *p*_*perm*_ = 0.070; Session *2, p*_*perm*_ < 0.001) but not in conditions where the implanted ear was primarily exposed to the distractor sound (select non-CI-side: Session 1, *p*_*perm*_ = 0.755; Session *2, p*_*perm*_ = 0.09; suppress CI-side: Session 1, *p*_*perm*_ = 0.956; Session *2, p*_*perm*_ = 0.298).

### Side of cochlear implant affects attentional modulation of alpha power

To test CI users’ neural signatures of auditory spatial attention to competing speech stimuli, we analysed the spatial attention-driven hemispheric lateralization of alpha oscillations (7–13 Hz) in the EEG. In agreement with a previous investigation with a similar paradigm in normal hearing listeners (Wöstmann et al., 2019), we found relatively higher posterior left-hemispheric alpha power following a leftward versus rightward spatial cue, and vice versa for right-hemispheric alpha power (Fig. 5A&B). Consistent with this, a repeated-measures ANOVA revealed a main effect of Hemisphere (left vs. right) on the alpha modulation index (AMI; *F*_1, 13_ = 6.451; *p* = 0.025) but no significant main effects of Session (1 vs. 2), CI status (on vs. off), or any interaction (all *p* > 0.2).

**Figure 5.**
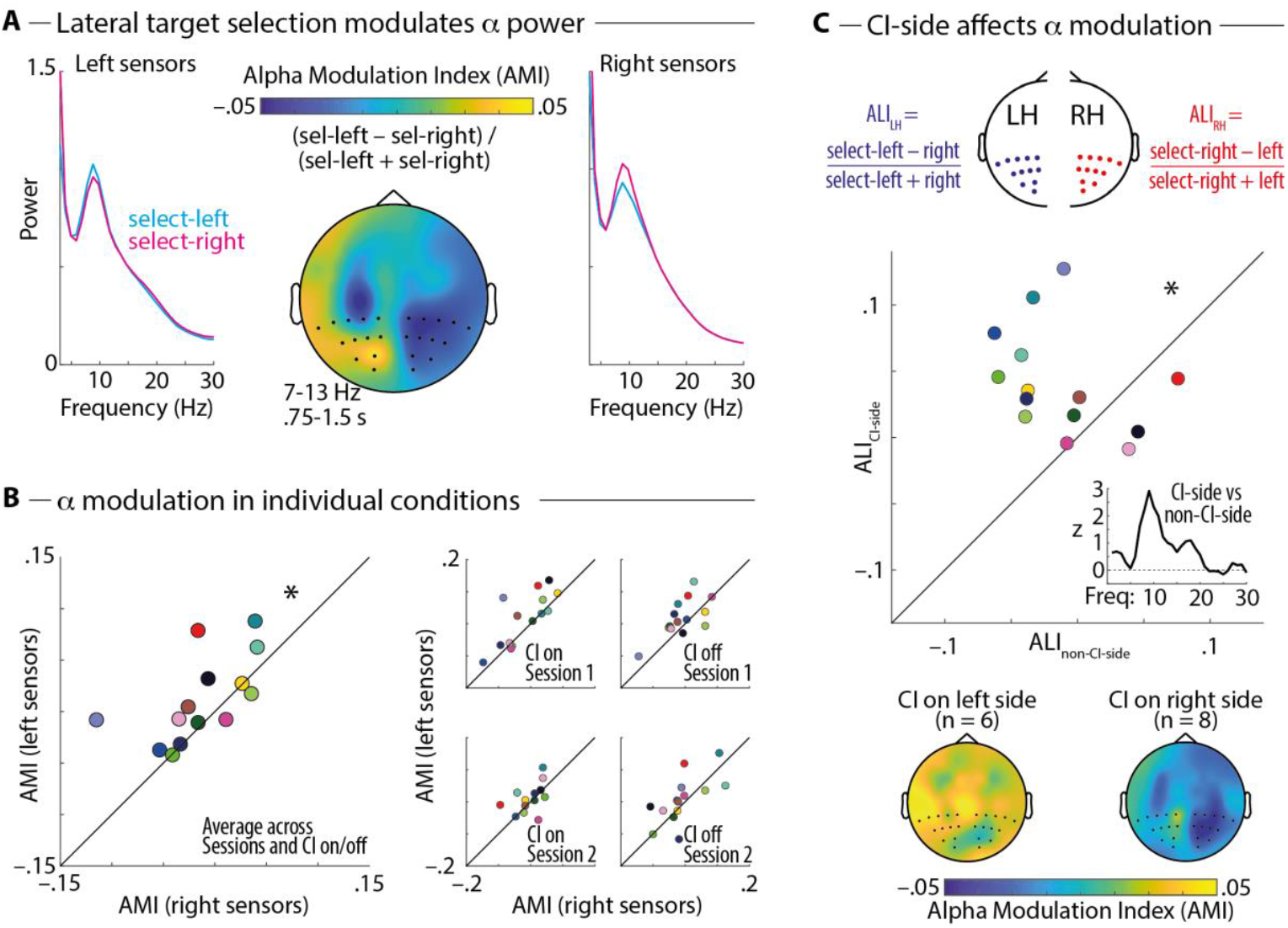
(**A**) Spectra show oscillatory power for conditions with the attended loudspeaker on the left or right side in the time interval in-between spatial cue presentation and stimulus onset, averaged across sessions (N = 18 in Session 1, N = 14 in Session 2) and CI status (on, off). The topographic map shows the attentional modulation index (AMI) in the alpha frequency band (7–13 Hz), calculated as (Power_select-left_ – Power_select-right_) / (Power_select-left_ + Power_select-right_). (**B**) 45-degree plots show AMI averaged across posterior right-(x-axis) versus left-hemispheric (y-axis) sensors for participants who participated in both sessions (N = 14 complete datasets). Note that datapoints above the diagonal indicate the expected pattern of higher AMI values at sensors on the left versus right hemisphere. * Significant main effect of Hemisphere (left vs. right) on AMI in repeated-measures ANOVA (*p* < 0.05). (**C**) Top: Illustration of the approach to calculate alpha lateralization indices (ALI) for the left (LH) and right (RH) hemispheres separately. Middle: 45-degree plot shows the ALI on the non-CI-side (x-axis) versus CI-side (y-axis). * Significant main effect of Hemisphere (CI-side vs. non-CI-side) on ALI in repeated-measures ANOVA (*p* < 0.05). Inset shows z-values for the contrast of lateralization indices on the CI-versus non-CI-side for individual frequencies (1-30 Hz). Bottom: Topographic maps show AMI separately for participants with the CI on the left versus right side.

Although we found before in a larger sample of normal hearing listeners (Wöstmann et al., 2019) a small but significant reversal of the alpha lateralization when contrasting trials with the spatial cue to the front and the distractor on the left versus right side (i.e., suppress-left vs. suppress-right conditions), this effect was not significant in the present dataset (*F*_1, 13_ = 0.894; *p* = 0.362).

In normal-hearing listeners, attentional modulation of lateralized alpha oscillations is typically assumed to be symmetrical between the two hemispheres. However, since spatial attention allocation in unilateral CI users is presumably affected by strong differences in hearing abilities on the left versus right side, we tested for asymmetries in hemispheric alpha lateralization. To this end, we calculated two alpha lateralization indices (ALI), one contrasting attentional selection of targets on the ipsi-versus contralateral side for electrodes on the left hemisphere, and another one for electrodes on the right hemisphere (Fig. 5C). A repeated-measures ANOVA revealed a main effect of Hemisphere (CI-side vs. non-CI-side), indicating that attentional alpha modulation was stronger in the hemisphere on the CI-side (*F*_1, 13_ = 9.045; *p* = 0.01). Critically, this effect was specific to the alpha frequency range (Fig. 5C, inset). This rules out the possibility that differences in EEG data preprocessing associated with the interpolation of missing electrodes on the CI-side might have introduced differential sensitivity for attentional modulation of oscillatory power between the hemispheres. Post-hoc tests revealed that alpha lateralization was significant in the hemisphere on the CI-side (*p*_*perm*_ < 0.001) but not in the hemisphere on the non-CI-side (*p*_*perm*_ = 0.276). There were no significant main effects of Session (1 vs. 2), CI status (on vs. off) or any interaction effects (all *p* > 0.37).

In a control analysis, we found that the effect of CI side on alpha modulation could not be explained by differential alpha modulation in the left versus right hemisphere independent of CI side (non-significant main effect of left vs. right hemisphere: *F*_1, 13_ = 0.094; *p* = 0.764). No clear pattern of correlations between behavioural and EEG measures in the present study could be found (see Fig. S3).

## Discussion

Here, we performed a longitudinal study to investigate electrophysiological indices of auditory temporal coding and spatial attention allocation in listeners who adapt to listening with a unilateral cochlear implant (CI). The main findings are as follows. First, six months post CI activation, the auditory steady-state response (ASSR) relatively decreased for 4-Hz amplitude-modulated sounds but increased for 40-Hz sounds. Second, spatial attention induced a robust lateralization of neural alpha oscillations in unilateral CI users, which was stable over time. Third, attentional alpha modulation was stronger in the hemisphere contralateral to the non-implanted ear, irrespective of time or of performing the task with the CI on versus off.

### Auditory perceptual processing following CI activation

Hearing with a CI has a profound impact on the sensory perception of acoustic signals. For instance, cochlear implantation is followed by increased metabolic activity in auditory cortex (especially contralateral; Lazeyras et al., 2002). Most CI users require several months to reach maximal perceptual performance (e.g., Glennon et al., 2020; Hamzavi et al., 2003). Restored hearing through CIs is associated with a broad range of plastic changes in the human brain (Giraud et al., 2002). In experienced adult CI users, auditory-induced cortical activity correlates with speech perception (K. M. J. Green et al., 2005). Here, we tested auditory phase-locking to amplitude-modulated sounds to help unveil the underlying neural adaptation to listening with a CI.

Vocoding has been used to simulate some aspects of hearing with a CI in normal-hearing listeners (e.g., Green et al., 2002). Using this method, we found previously that unilateral noise-vocoding delayed the attentional modulation of neural phase-locked responses to the envelope of speech stimuli (Kraus et al., 2021), which has a modulation spectrum peaking at ∼4 Hz (Ding et al., 2017). In the present study, however, we employed a passive listening task to probe the purely perceptual aspects of processing sound irrespective of attention focus.

Preferential tracking of acoustic stimuli in the theta (∼4–7 Hz) and gamma (∼30–50 Hz) ranges has been found before (Lakatos et al., 2005; Teng et al., 2017), and matches well with our observation of stronger phase-locking at 4 and 40 Hz compared with 20 Hz.

Differential modulation of 4-vs. 40-Hz ASSR during adaptation to a CI is presumably based on two underlying mechanisms. First, a relatively increased 40-Hz ASSR in the 7^th^ vs. 1^st^ month might suggest improved auditory temporal integration (Picton, 2013). Auditory temporal processing is important for speech processing and thus, for successful communication. Critically, unilateral CI users performed the passive listening task with both ears, that is, with the implanted and non-implanted ear. It is conceivable that better integration of electric hearing (through the CI) with acoustic hearing (on the non-implanted side) is associated with higher temporal coding acuity over time during adaptation to listening with the CI. Additionally, the increased 40-Hz ASSR might indicate increased inhibitory processing at the level of auditory cortical circuits (Toso et al., 2024).

Second, it might appear counterintuitive that the 4-Hz ASSR did not increase but decrease over time in the present study. However, larger amplitudes of cortical phase-locked responses to slow modulation frequencies have been observed before at older age (Decruy et al., 2019; Presacco et al., 2016) and in participants with hearing loss (Fuglsang et al., 2020; Orf et al., 2025). Mechanistically, a larger EEG amplitude might indicate a cortical overrepresentation, potentially compensating for weaker subcortical representations (Presacco et al., 2019). The observed decrease of the 4-Hz ASSR over time might thus indicate decreasing cortical overrepresentation and restoration of the excitation-inhibition balance. Future studies should test the hypothesis that individual differences in the balance of the 4-vs 40-Hz ASSR are explained by the duration of deafness before CI implantation in larger samples.

### Biased auditory spatial attention in unilateral CI users

The present study probed unilateral CI users’ speech reception in different close-to-real life arrangements of spatially competing target and distractor speech. In general agreement with previous research, we found better performance with versus without the CI (Távora-Vieira et al., 2015) and a tendency of this effect to increase over time (see also Dillon et al., 2020). Note that speech presentation levels were individually adapted to CI users’ hearing levels in both sessions (1^st^ and 7^th^ month), which might have reduced the ability to detect longitudinal effects on behavioral performance.

Importantly, previous research in normal-hearing listeners has shown that better acoustics (i.e., less degradation) not only has the intended effect of improved reception of target speech. Beter acoustics also favor increased interference by speech distractors (e.g., Ellermeier et al., 2015; Wöstmann et al., 2017; Wöstmann & Obleser, 2016). Based on this, one might have expected that hearing with a unilateral CI aids target speech reception on the CI side but at the same time increases interference effects of speech distractors.

To the contrary, our results demonstrate a benefit of listening with a unilateral CI in all conditions, especially (and statistically significant) in those where the target speech signal is primarily presented on the CI side. Thus, a unilateral CI does not uniquely benefit processing speech on the implanted side, but instead aids auditory object formation and speech perception more generally (see also Barchet et al., 2025; Chen et al., 2025; Fiedler et al., 2019).

At the neural level, in agreement with a previous study in bilateral CI users (Paul et al., 2020), we here demonstrate robust spatial attention-induced modulation of lateralized alpha oscillations in unilateral CI users. Alpha power relatively increased in the hemisphere ipsilateral to anticipated target sound and decreased in the contralateral hemisphere. Alpha lateralization was strongest at parietal electrode sites, which is commensurate with modulation in general attention networks rather than in auditory cortex regions (Banerjee et al., 2011). The reversal of alpha power lateralization for anticipated distractors on the left versus right side, which we found before in a larger sample of normal-hearing participants (Wöstmann et al., 2019), was not observed in CI users. This, however, does not necessarily imply reduced distractor suppression in unilateral CI users, since neural signatures of distractor suppression are typically smaller in size and thus require larger samples to be detected (Wöstmann et al., 2022).

Some prior work suggests reduced attentional alpha modulation in hearing-impaired compared with normal-hearing listeners (Bonacci et al., 2019). Although the present study did not include a normal-hearing control group, the size of the observed modulation of lateralized alpha power is comparable to a previous study with normal-hearing listeners that used a similar paradigm and the same EEG recording setup (Wöstmann et al., 2019). In sum, the present results provide an important proof of principle that hemispheric lateralization of alpha oscillations serves as a robust neural signature of auditory spatial attention deployment in CI users.

Typically, the hemispheric lateralization of alpha oscillations is assumed to be largely symmetrical between the left and right hemispheres. However, inter-individual variability of left-versus -right hemispheric attentional modulation of alpha power has been observed before and could partly be explained by asymmetry of subcortical structures (Ghafari et al., 2024; Mazzetti et al., 2019; Schulz & Wöstmann, 2024). Here, we substantially add to this line of research and demonstrate that asymmetric hearing in unilateral CI users is associated with stronger attentional modulation of alpha oscillations in the hemisphere contralateral to the non-implanted ear.

In theory, more pronounced spatial attention-driven alpha modulation in one hemisphere implies that the sensory input on the contralateral side is enhanced more when it is a target and/or suppressed more when it is a distractor (Schneider et al., 2021; Strauß et al., 2014). Our findings suggest that unilateral CI users apply auditory attentional processing predominantly in anticipation of auditory targets to the non-implanted ear (note that distractors were always presented in the front in target-lateral conditions). In unilateral CI users, task-specific effects of better-ear-listening have been found before (e.g., Williges et al., 2019), and it is likely that the non-implanted ear is the better ear for many participants in the present study. In general, our results highlight the importance of considering bilateral auditory input in CI rehabilitation (see also Chen et al., 2025).

Critically, the observed asymmetry of alpha lateralization was stable across time (7^th^ vs 1^st^ month post CI activation) and remained unmodulated by performing the task with versus without the CI. Absence of these effects could have been caused by our careful matching of task difficulty across conditions. That is, CI users’ performance was titrated to 50% correct in all conditions,meaning that task difficulty was in theory equal in the 1^st^ vs. 7^th^ month and with the CI on vs. off. Nevertheless, we tentatively conclude that the observed alpha asymmetry was driven by the history and presence of profound asymmetric hearing loss, rather than unilateral CI use. Future studies should test whether the asymmetric neural implementation of auditory spatial attention is present already before CI implantation and whether it declines after maximal performance in hearing with the unilateral CI is reached.

## Limitations

Of note, there are some limitations of the present study that should be considered. First, the sample size was small but the longitudinal nature of the study in part compensates for this lack in quantity by providing rich (i.e., “deep”) data from audiological tests, behavioral listening tasks, as well as EEG recordings in a passive listening test and in an active spatial attention task, in the 1^st^ and 7^th^ month following CI activation.

Second, the present study did not include a control group of listeners who did not receive a CI. Thus, it might be that some of the presumed CI-induced effects are partly driven by CI-independent learning mechanisms. While improving scores on audiological tests (FMST, SSQ, AMRD) might indeed be affected by learning, CI benefits in the spatial task were calculated by contrasting the CI-on with the CI-off condition in the 7^th^ versus 1^st^ month following CI activation. Thus, potential learning effects over time are being controlled for.

Third, CI conditions were tested in a fixed order (on, off) in the spatial listening task. In theory, the CI benefit might be somewhat overestimated and in part driven by adaptation to the task. However, we consider this rather unlikely since our calculation of speech reception thresholds omitted the first block for each experimental condition and thus focused on the later period where thresholds were stable.

## Conclusion

The electrophysiological evidence in our data suggests a longitudinal change in phase-locking to slower (4 Hz) versus faster (40 Hz) amplitude modulations in the first six months following activation of a unilateral CI. Irrespective of listening with or without the CI, unilateral CI users exhibit stronger neural attentional modulation associated with the non-implanted side, which emphasizes the importance of listening with both ears to unilateral CI users. These data provide first steps towards a mechanistic account of auditory perceptual and attentional adaptation to listening with a cochlear implant.

## Acknowledgements

This research was supported through a grant provided by Cochlear Research & Development Ltd under reference number IIR-2052 to MW, JO, and RS. We consulted generative AI (ChatGPT 4o, OpenAI Inc.) for help with minor text editing. As authors, we take full responsibility for these AI-mediated changes.

## References

Ahveninen, J., Huang, S., Belliveau, J. W., Chang, W.-T., & Hämäläinen, M. (2013). Dynamic Oscillatory Processes Governing Cued Orienting and Allocation of Auditory Attention. Journal of Cognitive Neuroscience, 25(11), 1926–1943. 10.1162/jocn_a_00452

Anderson, S., Roque, L., Gaskins, C. R., Gordon-Salant, S., & Goupell, M. J. (2020). Age-Related Compensation Mechanism Revealed in the Cortical Representation of Degraded Speech. Journal of the Association for Research in Otolaryngology, 21(4), 373–391. 10.1007/s10162-020-00753-4

Banerjee, S., Snyder, A. C., Molholm, S., & Foxe, J. J. (2011). Oscillatory Alpha-Band Mechanisms and the Deployment of Spatial Attention to Anticipated Auditory and Visual Target Locations: Supramodal or Sensory-Specific Control Mechanisms? Journal of Neuroscience, 31(27), Article 27. 10.1523/JNEUROSCI.4660-10.2011

Barchet, A. V., Bruera, A., Wend, J., Rimmele, J. M., Obleser, J., & Hartwigsen, G. (2025). Attentional engagement with target and distractor streams predicts speech comprehension in multitalker environments (p. 2025.04.04.647157). bioRxiv. 10.1101/2025.04.04.647157

Bonacci, L. M., Dai, L., & Shinn-Cunningham, B. G. (2019). Weak neural signatures of spatial selective auditory attention in hearing-impaired listeners. The Journal of the Acoustical Society of America, 146(4), 2577–2589. 10.1121/1.5129055

Bonnefond, M., & Jensen, O. (2024). The role of alpha oscillations in resisting distraction. Trends in Cognitive Sciences, 0(0). 10.1016/j.tics.2024.11.004

Brainard, D. H. (1997). The Psychophysics Toolbox. Spatial Vision, 10(4), 433–436. 10.1163/156856897X00357

Chen, Y.-P., Neff, P., Leske, S., Wong, D. D. E., Peter, N., Obleser, J., Kleinjung, T., Dimitrijevic, A., Dalal, S. S., & Weisz, N. (2025). Cochlear implantation in adults with acquired single-sided deafness improves cortical processing and comprehension of speech presented to the non-implanted ears: A longitudinal EEG study. Brain Communications, 7(1), fcaf001. 10.1093/braincomms/fcaf001

Cruz, G., Melcón, M., Sutandi, L., Palva, J. M., Palva, S., & Thut, G. (2025). Oscillatory Brain Activity in the Canonical Alpha-Band Conceals Distinct Mechanisms in Attention. Journal of Neuroscience, 45(1). 10.1523/JNEUROSCI.0918-24.2024

David, W., Verwaerde, E., Gransier, R., & Wouters, J. (2023). Effects of analysis window on 40-Hz auditory steady-state responses in cochlear implant users. Hearing Research, 438, 108882. 10.1016/j.heares.2023.108882

Decruy, L., Vanthornhout, J., & Francart, T. (2019). Evidence for enhanced neural tracking of the speech envelope underlying age-related speech-in-noise difficulties. Journal of Neurophysiology, 122(2), 601–615. 10.1152/jn.00687.2018

Deng, Y., Choi, I., & Shinn-Cunningham, B. (2020). Topographic specificity of alpha power during auditory spatial attention. NeuroImage, 207, 116360. 10.1016/j.neuroimage.2019.116360

Dillon, M. T., Buss, E., Rooth, M. A., King, E. R., McCarthy, S. A., Bucker, A. L., Deres, E. J., Richter, M. E., Thompson, N. J., Canfarotta, M. W., O’Connell, B. P., Pillsbury, H. C., & Brown, K. D. (2020). Cochlear Implantation in Cases of Asymmetric Hearing Loss: Subjective Benefit, Word Recognition, and Spatial Hearing. Trends in Hearing, 24, 2331216520945524. 10.1177/2331216520945524

Dimitrijevic, A., Smith, M. L., Kadis, D. S., & Moore, D. R. (2019). Neural indices of listening effort in noisy environments. Scientific Reports, 9(1), Article 1. 10.1038/s41598-019-47643-1

Dincer D’Alessandro, H., Ballantyne, D., Boyle, P. J., De Seta, E., DeVincentiis, M., & Mancini, P. (2018). Temporal Fine Structure Processing, Pitch, and Speech Perception in Adult Cochlear Implant Recipients. Ear and Hearing, 39(4), 679. 10.1097/AUD.0000000000000525

Ding, N., Patel, A. D., Chen, L., Butler, H., Luo, C., & Poeppel, D. (2017). Temporal modulations in speech and music. Neuroscience & Biobehavioral Reviews, 81, 181–187. 10.1016/j.neubiorev.2017.02.011

Ellermeier, W., Kattner, F., Ueda, K., Doumoto, K., & Nakajima, Y. (2015). Memory disruption by irrelevant noise-vocoded speech: Effects of native language and the number of frequency bands. The Journal of the Acoustical Society of America, 138(3), Article 3. 10.1121/1.4928954

Erb, J., Ludwig, A. A., Kunke, D., Fuchs, M., & Obleser, J. (2019). Temporal Sensitivity Measured Shortly After Cochlear Implantation Predicts 6-Month Speech Recognition Outcome. Ear and Hearing, 40(1), Article 1. 10.1097/AUD.0000000000000588

Eshraghi, A. A., Nazarian, R., Telischi, F. F., Rajguru, S. M., Truy, E., & Gupta, C. (2012). The Cochlear Implant: Historical Aspects and Future Prospects. The Anatomical Record, 295(11), 1967–1980. 10.1002/ar.22580

Farahani, E. D., Goossens, T., Wouters, J., & van Wieringen, A. (2017). Spatiotemporal reconstruction of auditory steady-state responses to acoustic amplitude modulations: Potential sources beyond the auditory pathway. NeuroImage, 148, 240–253. 10.1016/j.neuroimage.2017.01.032

Fiedler, L., Wöstmann, M., Herbst, S. K., & Obleser, J. (2019). Late cortical tracking of ignored speech facilitates neural selectivity in acoustically challenging conditions. NeuroImage, 186, 33–42. 10.1016/j.neuroimage.2018.10.057

Fuglsang, S. A., Märcher-Rørsted, J., Dau, T., & Hjortkjær, J. (2020). Effects of Sensorineural Hearing Loss on Cortical Synchronization to Competing Speech during Selective Attention. Journal of Neuroscience, 40(12), Article 12. 10.1523/JNEUROSCI.1936-19.2020

Galambos, R., Makeig, S., & Talmachoff, P. J. (1981). A 40-Hz auditory potential recorded from the human scalp. Proceedings of the National Academy of Sciences, 78(4), 2643–2647. 10.1073/pnas.78.4.2643

Gatehouse, S., & Noble, W. (2004). The Speech, Spatial and Qualities of Hearing Scale (SSQ). International Journal of Audiology, 43(2), 85–99. 10.1080/14992020400050014

Ghafari, T., Mazzetti, C., Garner, K., Gutteling, T., & Jensen, O. (2024). Modulation of alpha oscillations by attention is predicted by hemispheric asymmetry of subcortical regions. eLife, 12. 10.7554/eLife.91650.2

Giraud, A. L., Truy, E., & Frackowiak, R. (2002). Imaging Plasticity in Cochlear Implant Patients. Audiology and Neurotology, 6(6), 381–393. 10.1159/000046847

Glennon, E., Svirsky, M. A., & Froemke, R. C. (2020). Auditory cortical plasticity in cochlear implant users. Current Opinion in Neurobiology, 60, 108–114. 10.1016/j.conb.2019.11.003

Green, K. M. J., Julyan, P. J., Hastings, D. L., & Ramsden, R. T. (2005). Auditory cortical activation and speech perception in cochlear implant users: Effects of implant experience and duration of deafness. Hearing Research, 205(1), 184–192. 10.1016/j.heares.2005.03.016

Green, T., Faulkner, A., & Rosen, S. (2002). Spectral and temporal cues to pitch in noise-excited vocoder simulations of continuous-interleaved-sampling cochlear implants. The Journal of the Acoustical Society of America, 112(5), 2155–2164. 10.1121/1.1506688

Grent-’t-Jong, T., Brickwedde, M., Metzner, C., & Uhlhaas, P. J. (2023). 40-Hz Auditory Steady-State Responses in Schizophrenia: Toward a Mechanistic Biomarker for Circuit Dysfunctions and Early Detection and Diagnosis. Biological Psychiatry, 94(7), 550–560. 10.1016/j.biopsych.2023.03.026

Haegens, S., Händel, B. F., & Jensen, O. (2011). Top-Down Controlled Alpha Band Activity in Somatosensory Areas Determines Behavioral Performance in a Discrimination Task. Journal of Neuroscience, 31(14), 5197–5204. 10.1523/JNEUROSCI.5199-10.2011

Hamzavi, J., Baumgartner, W., Pok, S. M., Franz, P., & Gstoettner, W. (2003). Variables Affecting Speech Perception in Postlingually Deaf Adults Following Cochlear Implantation. Acta Oto-Laryngologica, 123(4), 493–498. 10.1080/0036554021000028120

Heng, J., Cantarero, G., Elhilali, M., & Limb, C. J. (2011). Impaired perception of temporal fine structure and musical timbre in cochlear implant users. Hearing Research, 280(1), 192–200. 10.1016/j.heares.2011.05.017

Jiwani, S., Doesburg, S. M., Papsin, B. C., & Gordon, K. A. (2021). Effects of long-term unilateral cochlear implant use on large-scale network synchronization in adolescents. Hearing Research, 409, 108308. 10.1016/j.heares.2021.108308

Kießling, J., Grugel, L., Meister, H., & Meis, M. (2011). Übertragung der Fragebögen SADL, ECHO und SSQ ins Deutsche und deren Evaluation. Z. Audiol, 50, 6–16.

Kraus, F., Tune, S., Ruhe, A., Obleser, J., & Wöstmann, M. (2021). Unilateral Acoustic Degradation Delays Attentional Separation of Competing Speech. Trends in Hearing, 25, 23312165211013242. 10.1177/23312165211013242

Kwon, J. S., O’Donnell, B. F., Wallenstein, G. V., Greene, R. W., Hirayasu, Y., Nestor, P. G., Hasselmo, M. E., Potts, G. F., Shenton, M. E., & McCarley, R. W. (1999). Gamma Frequency–Range Abnormalities to Auditory Stimulation in Schizophrenia. Archives of General Psychiatry, 56(11), 1001–1005. 10.1001/archpsyc.56.11.1001

Lakatos, P., Shah, A. S., Knuth, K. H., Ulbert, I., Karmos, G., & Schroeder, C. E. (2005). An oscillatory hierarchy controlling neuronal excitability and stimulus processing in the auditory cortex. Journal of Neurophysiology, 94(3), 1904–1911. 10.1152/jn.00263.2005

Lazeyras, F., Boëx, C., Sigrist, A., Seghier, M. L., Cosendai, G., Terrier, F., & Pelizzone, M. (2002). Functional MRI of Auditory Cortex Activated by Multisite Electrical Stimulation of the Cochlea. NeuroImage, 17(2), 1010–1017. 10.1006/nimg.2002.1240

Lenarz, M., Sönmez, H., Joseph, G., Büchner, A., & Lenarz, T. (2012). Long-Term Performance of Cochlear Implants in Postlingually Deafened Adults. Otolaryngology–Head and Neck Surgery, 147(1), 112–118. 10.1177/0194599812438041

Levitt, H. (1971). Transformed Up-Down Methods in Psychoacoustics. The Journal of the Acoustical Society of America, 49(2B), 467–477. 10.1121/1.1912375

Liégeois-Chauvel, C., Lorenzi, C., Trébuchon, A., Régis, J., & Chauvel, P. (2004). Temporal Envelope Processing in the Human Left and Right Auditory Cortices. Cerebral Cortex, 14(7), 731–740. 10.1093/cercor/bhh033

Lin, F. R., Niparko, J. K., & Ferrucci, L. (2011). Hearing Loss Prevalence in the United States. Archives of Internal Medicine, 171(20), 1851–1853. 10.1001/archinternmed.2011.506

Luke, R., De Vos, A., & Wouters, J. (2017). Source analysis of auditory steady-state responses in acoustic and electric hearing. NeuroImage, 147, 568–576. 10.1016/j.neuroimage.2016.11.023

Luke, R., Van Deun, L., Hofmann, M., van Wieringen, A., & Wouters, J. (2015). Assessing temporal modulation sensitivity using electrically evoked auditory steady state responses. Hearing Research, 324, 37–45. 10.1016/j.heares.2015.02.006

Mazzetti, C., Staudigl, T., Marshall, T. R., Zumer, J. M., Fallon, S. J., & Jensen, O. (2019). Hemispheric Asymmetry of Globus Pallidus Relates to Alpha Modulation in Reward-Related Attentional Tasks. Journal of Neuroscience, 39(46), 9221–9236. 10.1523/JNEUROSCI.0610-19.2019

Müller, N., & Weisz, N. (2012). Lateralized Auditory Cortical Alpha Band Activity and Interregional Connectivity Pattern Reflect Anticipation of Target Sounds. Cerebral Cortex, 22(7), Article 7. 10.1093/cercor/bhr232

Obleser, J., Wöstmann, M., Hellbernd, N., Wilsch, A., & Maess, B. (2012). Adverse Listening Conditions and Memory Load Drive a Common Alpha Oscillatory Network. Journal of Neuroscience, 32(36), Article 36. 10.1523/JNEUROSCI.4908-11.2012

Oldfield, R. C. (1971). The assessment and analysis of handedness: The Edinburgh inventory. Neuropsychologia, 9(1), Article 1. 10.1016/0028-3932(71)90067-4

Oostenveld, R., Fries, P., Maris, E., & Schoffelen, J.-M. (2010). FieldTrip: Open Source Software for Advanced Analysis of MEG, EEG, and Invasive Electrophysiological Data. Computational Intelligence and Neuroscience, 2011, e156869. 10.1155/2011/156869

Orf, M., Hannemann, R., & Obleser, J. (2025). Does amplitude compression help or hinder attentional neural speech tracking? Journal of Neuroscience. 10.1523/JNEUROSCI.0238-24.2024

Paul, B. T., Uzelac, M., Chan, E., & Dimitrijevic, A. (2020). Poor early cortical differentiation of speech predicts perceptual difficulties of severely hearing-impaired listeners in multi-talker environments. Scientific Reports, 10(1), 6141. 10.1038/s41598-020-63103-7

Picton, T. (2013). Hearing in Time: Evoked Potential Studies of Temporal Processing. Ear and Hearing, 34(4), 385. 10.1097/AUD.0b013e31827ada02

Presacco, A., Simon, J. Z., & Anderson, S. (2016). Evidence of degraded representation of speech in noise, in the aging midbrain and cortex. Journal of Neurophysiology, 116(5), Article 5. 10.1152/jn.00372.2016

Presacco, A., Simon, J. Z., & Anderson, S. (2019). Speech-in-noise representation in the aging midbrain and cortex: Effects of hearing loss. PLOS ONE, 14(3), e0213899. 10.1371/journal.pone.0213899

Purcell, D. W., John, S. M., Schneider, B. A., & Picton, T. W. (2004). Human temporal auditory acuity as assessed by envelope following responses. The Journal of the Acoustical Society of America, 116(6), 3581–3593. 10.1121/1.1798354

Rosen, S. (1992). Temporal information in speech: Acoustic, auditory and linguistic aspects. Philosophical Transactions of the Royal Society of London. Series B: Biological Sciences, 336(1278), 367–373. 10.1098/rstb.1992.0070

Roß, B., Picton, T. W., & Pantev, C. (2002). Temporal integration in the human auditory cortex as represented by the development of the steady-state magnetic field. Hearing Research, 165(1), 68–84. 10.1016/S0378-5955(02)00285-X

Roth, T. N., Hanebuth, D., & Probst, R. (2011). Prevalence of age-related hearing loss in Europe: A review. European Archives of Oto-Rhino-Laryngology, 268(8), 1101–1107. 10.1007/s00405-011-1597-8

Schneider, D., Herbst, S. K., Klatt, L.-I., & Wöstmann, M. (2021). Target enhancement or distractor suppression? Functionally distinct alpha oscillations form the basis of attention. The European Journal of Neuroscience. 10.1111/ejn.15309

Schulz, M., & Wöstmann, M. (2024). Time to get deep. eLife, 13, e100755. 10.7554/eLife.100755

Shannon, R. V., Zeng, F.-G., Kamath, V., Wygonski, J., & Ekelid, M. (1995). Speech Recognition with Primarily Temporal Cues. Science, 270(5234), Article 5234. 10.1126/science.270.5234.303

Shinn-Cunningham, B. G. (2008). Object-based auditory and visual attention. Trends in Cognitive Sciences, 12(5), 182–186. 10.1016/j.tics.2008.02.003

Strauß, A., Wöstmann, M., & Obleser, J. (2014). Cortical alpha oscillations as a tool for auditory selective inhibition. Frontiers in Human Neuroscience, 8. https://www.frontiersin.org/articles/10.3389/fnhum.2014.00350

Távora-Vieira, D., De Ceulaer, G., Govaerts, P. J., & Rajan, G. P. (2015). Cochlear Implantation Improves Localization Ability in Patients With Unilateral Deafness. Ear and Hearing, 36(3), e93. 10.1097/AUD.0000000000000130

Teng, X., Tian, X., Rowland, J., & Poeppel, D. (2017). Concurrent temporal channels for auditory processing: Oscillatory neural entrainment reveals segregation of function at different scales. PLOS Biology, 15(11), Article 11. 10.1371/journal.pbio.2000812

Toso, A., Wermuth, A. P., Arazi, A., Braun, A., Jong, T. G.-’t, Uhlhaas, P. J., & Donner, T. H. (2024). 40 Hz Steady-State Response in Human Auditory Cortex Is Shaped by Gabaergic Neuronal Inhibition. Journal of Neuroscience, 44(24). 10.1523/JNEUROSCI.2029-23.2024

Williges, B., Wesarg, T., Jung, L., Geven, L. I., Radeloff, A., & Jürgens, T. (2019). Spatial Speech-in-Noise Performance in Bimodal and Single-Sided Deaf Cochlear Implant Users. Trends in Hearing, 23, 2331216519858311. 10.1177/2331216519858311

Wilson, T. W., Rojas, D. C., Reite, M. L., Teale, P. D., & Rogers, S. J. (2007). Children and Adolescents with Autism Exhibit Reduced MEG Steady-State Gamma Responses. Biological Psychiatry, 62(3), 192–197. 10.1016/j.biopsych.2006.07.002

Worden, M. S., Foxe, J. J., Wang, N., & Simpson, G. V. (2000). Anticipatory Biasing of Visuospatial Attention Indexed by Retinotopically Specific α-Bank Electroencephalography Increases over Occipital Cortex. Journal of Neuroscience, 20(6), Article 6. 10.1523/JNEUROSCI.20-06-j0002.2000

World Health Organization (Ed.). (2021). World report on hearing. https://www.who.int/publications/i/item/9789240020481

Wöstmann, M., Alavash, M., & Obleser, J. (2019). Alpha oscillations in the human brain implement distractor suppression independent of target selection. The Journal of Neuroscience, 39(49), Article 49. 10.1523/JNEUROSCI.1954-19.2019

Wöstmann, M., Herrmann, B., Maess, B., & Obleser, J. (2016). Spatiotemporal dynamics of auditory attention synchronize with speech. Proceedings of the National Academy of Sciences, 113(14), Article 14. 10.1073/pnas.1523357113

Wöstmann, M., Lim, S.-J., & Obleser, J. (2017). The Human Neural Alpha Response to Speech is a Proxy of Attentional Control. Cerebral Cortex, 27(6), Article 6. 10.1093/cercor/bhx074

Wöstmann, M., & Obleser, J. (2016). Acoustic Detail But Not Predictability of Task-Irrelevant Speech Disrupts Working Memory. Frontiers in Human Neuroscience, 538. 10.3389/fnhum.2016.00538

Wöstmann, M., Störmer, V. S., Obleser, J., Addleman, D. A., Andersen, Søren K., Gaspelin, N., Geng, J. J., Luck, S. J., Noonan, M. P., Slagter, H. A., & Theeuwes, J. (2022). Ten simple rules to study distractor suppression. Progress in Neurobiology, 213, 102269. 10.1016/j.pneurobio.2022.102269

Yang, X., Fiebelkorn, I. C., Jensen, O., Knight, R. T., & Kastner, S. (2024). Differential neural mechanisms underlie cortical gating of visual spatial attention mediated by alpha-band oscillations. Proceedings of the National Academy of Sciences, 121(45), e2313304121. 10.1073/pnas.2313304121

