## Supplementary Materials for "Neural Alpha Oscillations and Auditory Steady State Responses During Adaptation to a Cochlear Implant"

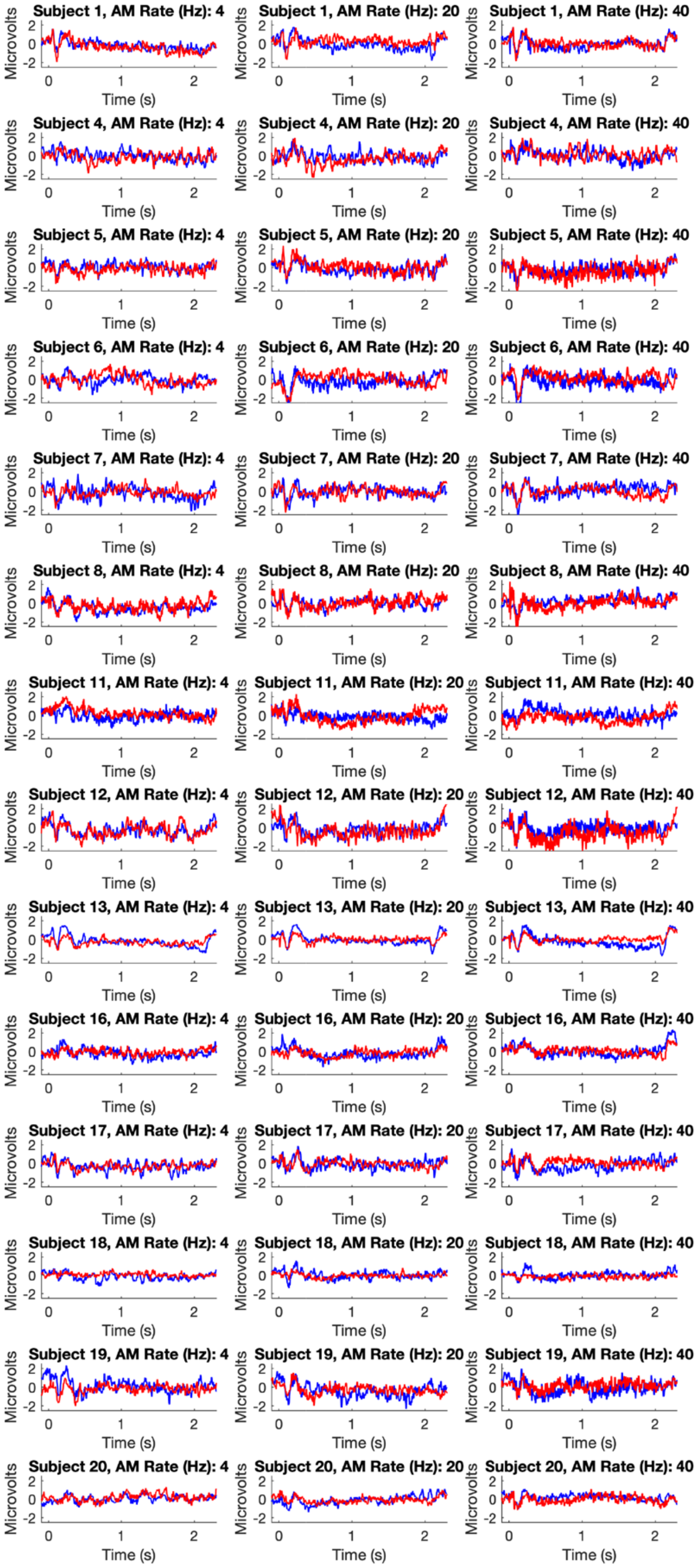


**Figure S1**. Single-subject event-related potential (ERP) to amplitude-modulated (AM) sounds (blue: session 1, red: session 2). The ERP has been averaged across 9 fronto-central channels and is shown for participants who completed both experimental sessions.

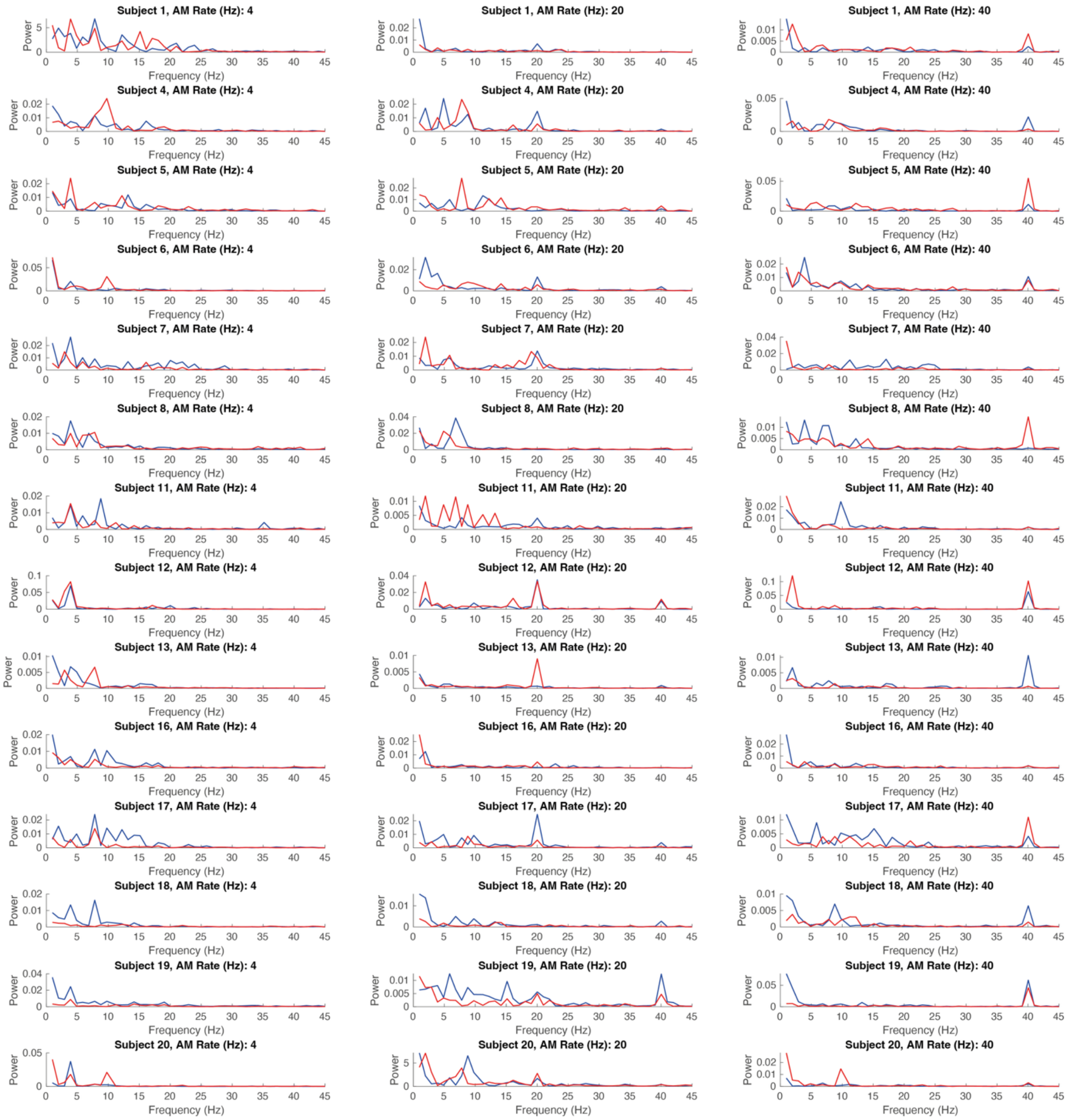


**Figure S2**. Single-subject spectral power, calculated on the ERP to amplitude-modulated (AM) sounds in the time-interval 0.5–2s, averaged across 9 fronto-central channels (blue: session 1, red: session 2). Spectra are shown for participants who completed both experimental sessions.

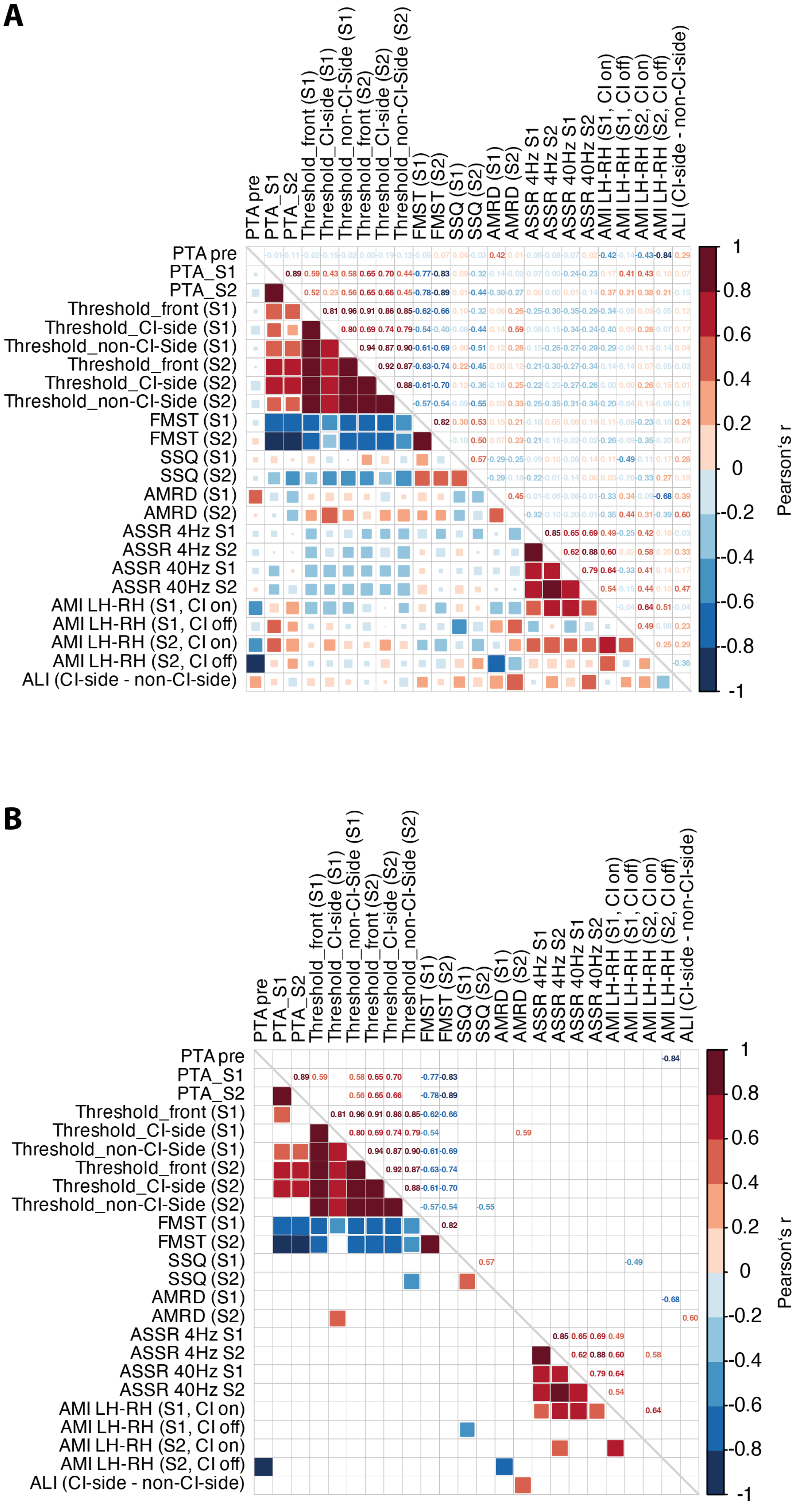


**Figure S3**. (**A**) Correlation heatmap for all variables of interest. (**B**) Same as A with all non-significant correlations ($p > .05$) removed. S1 = Session 1; S2 = Session 2. Note that due to the limited sample size (S1: N = 18; S2: N = 14), all correlations should be interpreted with caution.
